# SARS-CoV-2 Spike protein co-opts VEGF-A/Neuropilin-1 receptor signaling to induce analgesia

**DOI:** 10.1101/2020.07.17.209288

**Authors:** Aubin Moutal, Laurent F. Martin, Lisa Boinon, Kimberly Gomez, Dongzhi Ran, Yuan Zhou, Harrison J. Stratton, Song Cai, Shizhen Luo, Kerry Beth Gonzalez, Samantha Perez-Miller, Amol Patwardhan, Mohab M. Ibrahim, Rajesh Khanna

**Author notes:** Correspondence to: Dr. Rajesh Khanna, Department of Pharmacology, College of Medicine, University of Arizona, 1501 North Campbell Drive, P.O. Box 245050, Tucson, AZ 85724, USA. Contributed equally (Laurent F. Martin, Lisa Boinon, Kimberly Gomez, Dongzhi Ran, Yuan Zhou).

## Abstract

Global spread of severe acute respiratory syndrome coronavirus 2 (SARS-CoV-2) continues unabated. Binding of SARS-CoV-2’s Spike protein to host angiotensin converting enzyme 2 triggers viral entry, but other proteins may participate, including neuropilin-1 receptor (NRP-1). As both Spike protein and vascular endothelial growth factor-A (VEGF-A) – a pro-nociceptive and angiogenic factor, bind NRP-1, we tested if Spike could block VEGF-A/NRP-1 signaling. VEGF-A–triggered sensory neuronal firing was blocked by Spike protein and NRP-1 inhibitor EG00229. Pro-nociceptive behaviors of VEGF-A were similarly blocked via suppression of spontaneous spinal synaptic activity and reduction of electrogenic currents in sensory neurons. Remarkably, preventing VEGF-A/NRP-1 signaling was antiallodynic in a neuropathic pain model. A ‘silencing’ of pain via subversion of VEGF-A/NRP-1 signaling may underlie increased disease transmission in asymptomatic individuals.

## 1. Introduction

Severe acute respiratory syndrome coronavirus 2 (SARS-CoV-2) is the causative agent of COVID-19, a coronavirus disease that, as of August 24, has infected more than 23.5 million people and caused nearly 810,000 deaths worldwide [15]. Most patients infected with SARS-CoV-2 report mild to severe respiratory illness with symptoms such as fever, cough and shortness of breath [30]. On the other hand, a subset of patients who are diagnosed by a positive nuclei acids test but are either asymptomatic or minimally symptomatic [30]. Increasing evidence shows that asymptomatic individuals can spread the virus efficiently, and the emergence of these silent spreaders of SARS-CoV-2 has limited control of the pandemic [14; 41]. Pain is a rising concern in symptomatic patients, likely emanating from a direct attack of SARS-CoV-2 on cells and the “cytokine storm” unleashed by affected cells [51; 68]. Whether asymptomatic or minimally symptomatic individuals have reduced pain thresholds, or whether their pain is silenced is unknown, but either could contribute to increased disease transmission dynamics.

The surface expressed angiotensin converting enzyme 2 (ACE2) has been lionized as the main receptor for uptake of SARS-CoV-2 [22; 60; 64]. Emerging evidence points to a subset of ACE2 expressing sensory neurons [48] that synapse with spinal and brainstem CNS neurons to produce neurological effects, including headache and nerve pain [32; 34]. Curiously, ACE2 is not present in most neurons [48], despite increasing reports of neurological symptoms being common in COVID-19 patients [32]. Paradoxically, though the levels of ACE2 expression decline in aging [49], increased COVID-19 severity was noted in older patient populations, such as that of Italy’s [2], supporting the contention that ACE2 is not the sole gateway for entry of SARS-CoV-2 [1].

Two recent reports demonstrated that the SARS-CoV-2 Spike protein can bind to the b1b2 domain of the neuropilin-1 receptor (NRP-1). This interaction occurs through a polybasic amino acid sequence (^682^RRAR^685^), not conserved in SARS and MERS, termed the ‘C-end rule’ (CendR) motif, which significantly potentiates its entry into cells [6; 11]. Importantly, ‘omic’ analyses revealed a significant upregulation of *NRP-1* in biological samples from COVID-19 patients compared to healthy controls [6]. Using vascular endothelial growth factor-A (VEGF-A), a physiological ligand for the b1b2 pocket in NRP-1, we interrogated whether the Spike protein, the major surface antigen of SARS-CoV-2, could block VEGF-A/NRP-1 signaling to affect pain behaviors. Given parallels between the pro-nociceptive effects of VEGF-A in rodents [4; 58] and humans [23; 58] and clinical findings demonstrating increased VEGF-A levels in bronchial alveolar lavage fluid from COVID-19 patients [47] coupled with substantially lower levels in the sera of asymptomatic individuals compared to symptomatic patients [30], a secondary question was to test whether Spike protein could confer analgesia. We found that VEGF-A sensitizes nociceptor activity – a hallmark of neuropathic pain [59], which was blocked by the Spike protein and NRP-1 inhibitor EG00229 [26]. Furthermore, we identify a novel analgesic role for Spike protein, which is mirrored by NRP-1 inhibition.

## 2. Materials and Methods

An expanded version of all Methods is available in the **Supplementary Information** section.

### 2.1. Animals

Pathogen-free adult male and female Sprague-Dawley rats (225-250g; Envigo, Indianapolis, IN) were housed in temperature-controlled (23±3 °C) and light-controlled (12-h light/12-h dark cycle; lights on 07:00-19:00) rooms with standard rodent chow and water available ad libitum. The Institutional Animal Care and Use Committee of the College of Medicine at the University of Arizona approved all experiments. All procedures were conducted in accordance with the Guide for Care and Use of Laboratory Animals published by the National Institutes of Health. Animals were randomly assigned to treatment or control groups for the behavioral experiments. Animals were initially housed 3 per cage but individually housed after the intrathecal cannulation. All behavioral experiments were performed by experimenters who were blinded to the experimental groups and treatments.

### 2.2. ELISA□based NRP1–Spike protein binding assay

Plates (96□well, Nunc Maxisorp; Thermo Fisher Scientific, Waltham, MA, USA) were coated with human Neuropilin-1-Fc (10 ng per well, Cat# 50-101-8343, Fisher, Hampton, NH) and incubated at room temperature overnight. The following day, the plates were washed and blocked with 3% BSA in PBS to minimize non□specific adsorptive binding to the plates. SARS-CoV-2 Spike protein (S1 domain aa16-685, Cat# Z03485, Genscript, Piscataway, NJ) was added at concentrations ranging from 0.07 to 500 to nM. As a negative control, some wells received PBS containing 3% BSA. The plates were incubated at room temperature with shaking for 3 h. Next, the plates were washed with PBS to eliminate unbound protein. Bound SARS-CoV-2 Spike was detected by antiLJHis probe HRP (Cat#15165; Thermo Fisher Scientific). Some wells received no antibody (no antibody control). Tetramethylbenzidine (Cat#DY999, R&D Systems, St. Louis, MO) was used as the colorimetric substrate. The optical density of each well was determined immediately, using a microplate reader (Multiskan Ascent; Thermo Fisher Scientific) set to 450 nm with a correction wavelength of 570 nm. Data were analyzed by non-linear regression analysis using GraphPad Prism 8 (GraphPad, San Diego, CA, USA).

### 2.3. Preparation of acutely dissociated dorsal root ganglion neurons

Dorsal root ganglia from all levels were acutely dissociated using methods as described previously [18]. Rat DRG neurons were isolated from 100g female Sprague-Dawley rats using previously developed procedures [36].

### 2.4. Culturing primary dorsal root ganglia (DRG) neurons and micro-electrode array (MEA) analysis

Dissociated DRG neurons were maintained in media containing Neurobasal (Cat# 21103049, Thermofisher), 2% B-27 (Cat# 17504044, Thermofisher), 1% penicillin/streptomycin sulfate from 10,000 µg/ml stock, 30 ng/ml nerve growth factor, and 10% fetal bovine serum (Hyclone). Collected cells were re-suspended in DRG media and seeded as a 10-µl drop on the poly-D-lysine coated electrodes of the micro-electrode array (MEA) (24-well plate, Cat# MED-Q2430L, MED64, Japan). The cells were allowed to adhere for 30 min and then flooded with DRG media. The next day, cells were analyzed on a MED64 presto where 24 wells, with each containing 16 electrodes, could be recorded simultaneously. Cells were treated with the indicated neuropilin 1 ligands Semaphorin 3A (3nM; Cat#5926-S3-025, R&D Systems), VEGF-A_165_ (1nM, Cat#P4853, Abnova) or VEGF-B (1nM, Cat#RPU44324, Biomatik) for 30 min before recording. Treatments with either Spike (100nM, Cat#Z03485, Genscript) or the NRP-1 inhibitor EG-00229 (30µM, Cat#6986, Tocris) was done 30min before adding VEGF-A. The data was analyzed on MEA symphony and Mobius offline toolkit to extract the firing rate of the active electrodes before and after prolactin treatment. Firing rate is shown as Hz (event per second) for the electrodes that showed spontaneous activity.

### 2.5. Whole-cell patch recordings of Na^+^ and N-type (CaV2.2) Ca^2+^ currents in acutely dissociated DRG neurons

Recordings were obtained from acutely dissociated DRG neurons as described previously [25; 37]. The solutions and protocols for measuring sodium currents was exactly as descried [35]. The protocol for isolating N-type calcium currents was previously described by Khanna et al. [10; 28]. To isolate the N-type channel in DRGs, contributions of the high-and low-voltage-activated low calcium channel subtypes was blocked with the following subunit-selective blockers (all purchased from Alomone Labs, Jerusalem): Nifedipine (10 μM, L-type); ω-agatoxin GIVA (200 nM, P/Q-type) [33]; SNX-482 (200 nM, R-type) [39]; and TTA-P2 (1 µM, T-type)[10]. DRG neurons were interrogated with current-voltage (I-V) and activation/inactivation voltage protocols as previously described [16; 18; 35].

To determine the effect of VEGF-A application on voltage gated sodium and calcium currents, we incubated recombinant rat VEGF-A (1 nM in PBS) with DRG neurons for 30 minutes before whole-cell recordings. Additionally, recombinant Spike protein (100 nM in PBS) and the Neuropilin 1 (NRP-1) blocker EG00229 (30 μM in DMSO, Cat. No. 6986, Tocris Bioscience) were also applied to the culture medium for 30 minutes before recording. For experiments where the proteins were tested in combination, the Spike protein was added first for 30 minutes, followed by VEGF-A and EG00229 for another 30 minutes before recording commenced. The control conditions used either PBS, DMSO, or both to match the solutions used in the experimental conditions. The proteins and blocker were included at the same concentrations in the extracellular recording solution during all data acquisition.

### 2.6. Preparation of spinal cord slices

As described before [65], transverse 350-μm thick slices were prepared from young rats (postnatal 10-14 days) for electrophysiological recordings at RT. VEGFA (1nM), NRP-1 inhibitor (EG00229, 30 µM) and Spike protein were added directly to the recording solution as indicated. Slices were treated for 30 min prior to the recordings.

### 2.7. Electrophysiological recording in spinal cord slices by whole-cell patch clamp

*Substantia gelatinosa* neurons were recorded from using methods exactly as described previously [66].

### 2.8. Hind paw injection procedures

PBS vehicle (NaCl 137 mM, KCl 2.5 mM, Na_2_HPO_4_ 10 mM and KH_2_PO_4_ 1.8 mM), VEGF-A_165_

(10 nM), Spike (100 nM) and EG00229 (30 µM) were injected subcutaneously, alone or in combination, in the dorsum of the left hind paw. Rats were gently restrained under a fabric cloth, and 50 µL were injected using 0.5 mL syringes (27-G needles).

### 2.9. Implantation of intrathecal catheter

For intrathecal (i.t.) drug administration, rats were chronically implanted with catheters as described by Yaksh and Rudy [63].

### 2.10. Testing of allodynia

The assessment of tactile allodynia (i.e., a decreased threshold to paw withdrawal after probing with normally innocuous mechanical stimuli) using a series of calibrated fine (von Frey) filaments was done as described in Chaplan et al [8].

### 2.11. Spared nerve injury (SNI)

Nerve injury was inflicted as described previously [12]. Rats were tested on day 10 following the surgery, at which time they had developed stable and maximal mechanical allodynia.

### 2.12. Synapse enrichment and fractionation

Adult rats were anesthetized using isoflurane and decapitated. Spinal cords were removed, and the dorsal horn of the spinal cord was dissected as this structure contains the synapses arising from the DRG. Synaptosomes isolation was done as described previously [42]. Integrity of non-postsynaptic density (non-PSD) and PSD fractions was verified by immunoblotting. PSD95 was enriched in the PSD fractions while synaptophysin was enriched in non-PSD fractions (not shown). BCA protein assay was used to determine protein concentrations.

### 2.13. Immunoblot preparation and analysis

Protein concentrations were determined using the BCA protein assay (Cat# PI23225, Thermo Fisher Scientific, Waltham, MA). Indicated samples were loaded on 4-20% Novex® gels (Cat# EC60285BOX, Thermo Fisher Scientific, Waltham, MA). Proteins were transferred for 1h at 120 V using TGS (25mM Tris pH=8.5, 192mM glycine, 0.1% (mass/vol) SDS), 20% (vol/vol) methanol as transfer buffer to polyvinylidene difluoride (PVDF) membranes 0.45μm (Cat# IPVH00010, Millipore, Billerica, MA), pre-activated in pure methanol. After transfer, the membranes were blocked at room temperature for 1 hour with TBST (50 mM Tris-HCl, pH 7.4, 150 mM NaCl, 0.1 % Tween 20), 5% (mass/vol) non-fat dry milk, then incubated separately in the primary antibodies VEGFR2 (Cat#PA5-16487, ThermoFisher), pY1175 VEGFR2 (Cat#PA5-105167, ThermoFisher), Flotilin (Cat#F1180, Sigma) or Neuropilin-1 (Cat#sc-5307, Santa Cruz biotechnology) in TBST, 5% (mass/vol) BSA, overnight at 4°C. Following incubation in horseradish peroxidase-conjugated secondary antibodies from Jackson immunoresearch, blots were revealed by enhanced luminescence (WBKLS0500, Millipore, Billerica, MA) before exposure to photographic film. Films were scanned, digitized, and quantified using Un-Scan-It gel version 7.1 scanning software by Silk Scientific Inc.

### 2.14. Docking of VEGF-A Spike protein and EG00229 to NRP-1

Peptide from C-terminus of furin cleaved SARS-CoV-2 Spike protein 681-PRRAR-685 (blue sticks) was docked to NRP-1-b1 domain (white surface with binding site in red; PDB 6fmc [46]) using Glide (Schrödinger [19]). Molecular figures generated with PyMol 1.8 (Schrödinger, LLC.)

### 2.15. Statistical Analysis

All data was first tested for a Gaussian distribution using a D’Agostino-Pearson test (Prism 8 Software, Graphpad, San Diego, CA). The statistical significance of differences between means was determined by a parametric ANOVA followed by Tukey’s post hoc or a non-parametric Kruskal Wallis test followed by Dunn’s post-hoc test depending on if datasets achieved normality. Behavioral data with a time course were analyzed by Two-way ANOVA with Sidak’s post hoc test. Differences were considered significant if p≤ 0.05. Error bars in the graphs represent mean ± SEM. All data were plotted in Prism 8.

## 3. Results

### 3.1. Ligand specific engagement of NRP-1 signaling induces nociceptor activity and pain

Initially, we assessed the involvement of Spike and NRP-1 in the VEGF-A/NRP-1 pathway. An interaction between Spike (S1 domain aa 16-685, containing the CendR motif ^682^RRAR^685^) and the extracellular portion of NRP-1 was confirmed by enzyme-linked immunosorbent assay (ELISA) (Fig. 1A). We calculated an equilibrium constant of dissociation (Kd) for this interaction to be ∼166.2 nM (Fig. 1A). Next, we plated sensory neurons on multiwell microelectrode arrays (MEAs), an approach enabling multiplexed measurements of spontaneous, as well as stimulus-evoked extracellular action potentials from large populations of cells [9]. VEGF-A increased spontaneous firing of dorsal root ganglion (DRG) neurons, which was blocked by the S1 domain of the Spike protein and by the NRP-1 inhibitor EG00229 (Fig. 1A). In contrast, ligands VEGF-B (ligand for VEGFR1 – a co-receptor for NRP-1 [31]) and semaphorin 3A (Sema3A, ligand for plexin receptor – also a co-receptor for NRP-1) [13; 53]) did not affect the spontaneous firing of nociceptors (Fig. 1B, C). The lack of effect of VEGF-B and Sema3A rule out a role for VEGF-R1 and plexin, respectively, thus implicating a novel ligand-, VEGF-A, and receptor-, NRP-1, specific pathway driving nociceptor firing (Fig. 1D).

**Figure 1.**
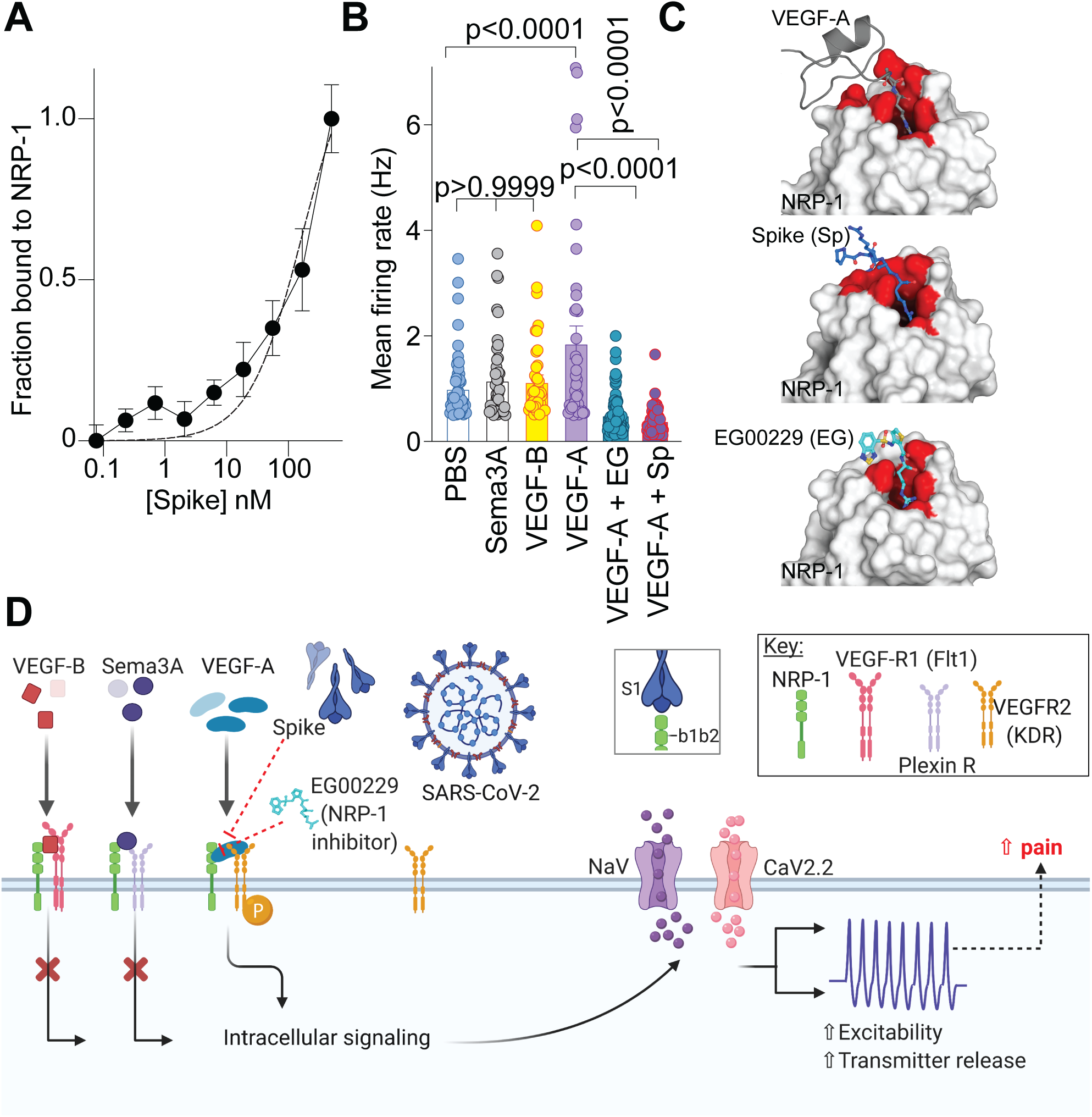
Ligand specific engagement of NRP-1 signaling induces nociceptor activity, promoting a pain-like phenotype. (A) Graph showing normalized NRP-1 binding to increasing concentrations of recombinant Spike protein (n=8 replicates per concentration). Data were normalized to wells with no antibody and background subtracted. The data was fit assuming a one site of binding mode and yielded a Kd of ∼166.2 nM. (**B**) Mean action potential firing rates (Hz, event per second) of cultured DRG sensory neurons incubated for 30 min with VEGF-B (3 nM), Sema3A (100 ng), VEGF-A (1 nM), VEGF-A plus Spike (100 nM) or VEGF-A plus NRP-1 inhibitor EG00229 (30 μM) [26]. Of the ligands tested, only VEGF-A, acting on VEGFR2, is a ligand for NRP-1 that triggers an increase in spontaneous firing of nociceptors. Data is shown as mean ± s.e.m. and was analyzed by non-parametric two-way analysis of variance (post hoc: Sidak). P values, versus control (PBS) or VEGF-A, are indicated. EG00229 is an NRP-1 inhibitor (PDB 3i97). (**C**) *Top left*: VEGF-A heparin binding domain (gray cartoon with R323 in sticks) in complex with the NRP-1 b1 domain (white surface with binding site in red; PDB 4deq [43]). *Top right*: Peptide from C-terminus of furin cleaved SARS-CoV-2 Spike protein 681-PRRAR-685 (blue sticks) docked to NRP1-b1 domain (white surface with binding site in red; PDB 6fmc [46]) using Glide (Schrödinger). Additional Spike residues 678-TNS-680 modeled for illustration purposes only (blue cartoon). *Bottom*: Compound EG00229 (cyan sticks) in complex with NRP-1 b1 domain ((white surface with binding site in red; PDB 3i97 [26]). (**D**) Schematic illustration of the hypothesis that SARS-CoV-2 Spike protein binding to NRP-1 b1b2 domain triggers an intracellular cascade that increases sodium and calcium channel activity to increase nociceptor activity culminating in enhanced pain.

As both VEGF-A and Spike protein share a common binding pocket on NRP-1 (Fig. 1C) [6; 11; 43], we asked if the Spike protein could block VEGF-A/NRP-1 signaling to affect pain behaviors. Consistent with previous reports [4; 58], we confirmed that VEGF-A is pro-nociceptive as intra-plantar injection of VEGF-A decreased both paw withdrawal thresholds (Fig. 2A, B and Table S1) and latencies to a thermal stimulus (Fig. 2C, D and Table S1) in male rats. Similar results were obtained in female rats as well (Fig. 2E-H and Table S1). Preventing VEGF-A from binding to NRP-1 with the NRP-1 inhibitor EG00229 or Spike from activating VEGF-A/NRP-1 signaling, blunted the mechanical allodynia and thermal hyperalgesia induced by VEGF-A alone (Fig. 2 and Table S1). Neither Spike nor EG00229 alone had any effect on these behaviors (Fig. 2 and Table S1) in either sex. Together, these data provide functional evidence that VEGF-A/NRP-1 signaling promotes a pain-like phenotype by sensitizing nociceptor activity (Fig. 1D).

**Figure 2.**
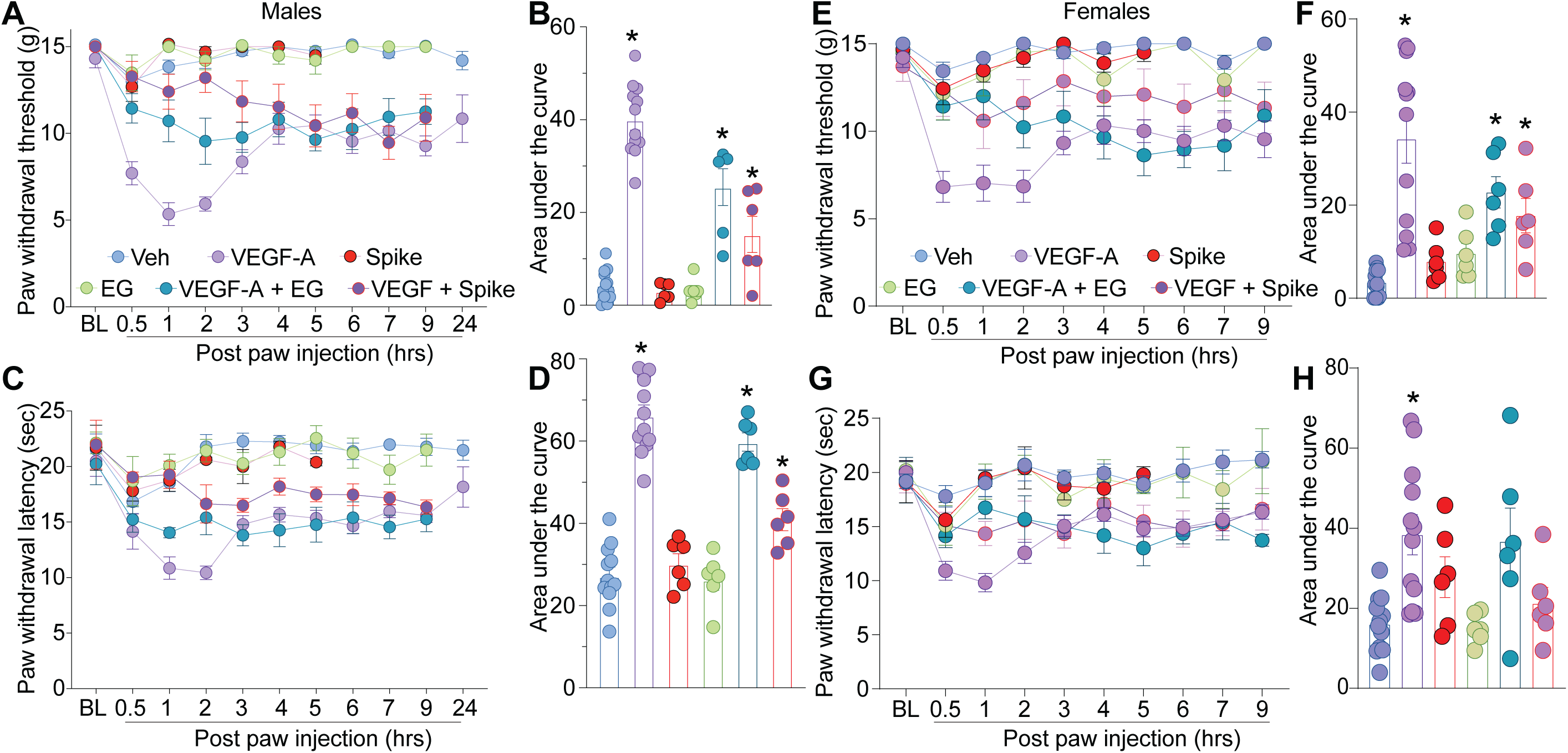
VEGF-A promotes a pain-like phenotype that is blocked by Spike protein or NRP-1 inhibition in male and female rats. Paw withdrawal thresholds (**A, B** – male and **E, F** – female) or latencies (**C, D** – male and **G, H** – female) for male and female naïve rats injected in the paw with VEGF-A (10 nM), Spike (100 nM), EG00229 (30 µM) or PBS (vehicle), alone or in combination (50 µl/rat; = 6-12). For clarity, statistical significance is not presented in the time course graphs, instead it is presented in Table S1. Panels **B, F, D** and **H** are the area under the curve for 0-24 hours. Data is shown as mean ± s.e.m. and was analyzed by non-parametric two-way analysis of variance where time was the within subject factor and treatment was the between subject factor (post hoc: Sidak), *p<0.05. Areas under the curve were compared by a one-way analysis of variance with Kruskal-Wallis post-hoc test. The experiments were analyzed by an investigator blinded to the treatment. For full statistical analyses see Table S1.

### 3.2. VEGF-A–mediated increases in DRG ion channel currents are normalized by disruption of VEGF-A/NRP-1 signaling

To gain insight into the mechanism by which VEGF-A contributed to increased nociceptor activity, we postulated that ion channels in DRGs may be affected, as these contribute to nociceptive plasticity [61]. Typical families of Na^+^ currents from small diameter DRG neurons are shown in Figure 3A. VEGF-A facilitated a 1.9–fold increase in total Na^+^ currents compared to vehicle (PBS)-treated DRGs, which was completely blocked by Spike protein (Fig. 3B, C). Spike protein alone did not affect Na^+^ currents (Fig. 3B, C and Table S1). Since this decreased current could arise from changes in channel gating, we determined if activation and inactivation kinetics of DRG Na^+^ currents were affected. Half-maximal activation and inactivation (V_1/2_), as well as slope values (*k*) for activation and inactivation, were no different between the conditions tested (Fig. 3D, E and Tables S1, S2), except for an ∼8 mV hyperpolarizing shift in sodium channel inactivation induced by co-treatment of VEGF-A and EG00229 (Table S2). Similar results were obtained for the NRP-1 inhibitor EG00229, which also inhibited the VEGF-A mediated increase in total Na^+^ currents (Fig. 3F-H and Table S1) but had no effect on the biophysical properties (Fig. 3I, J and Tables S1, S2).

**Figure 3.**
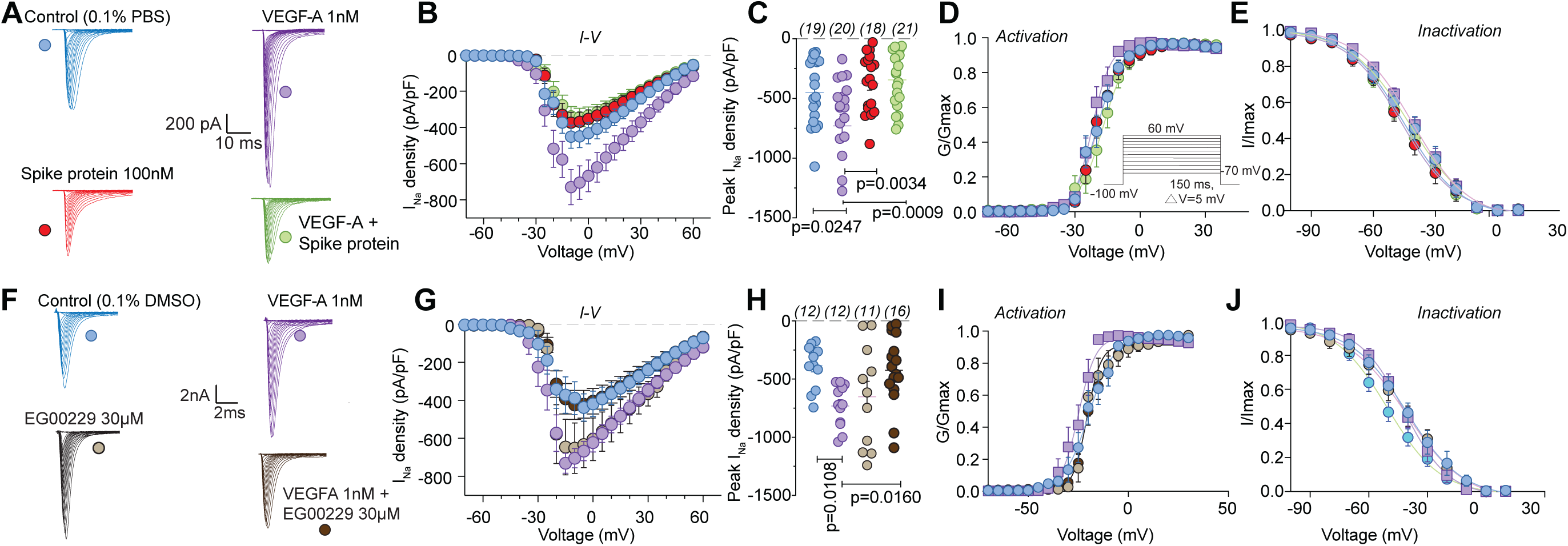
VEGF-A–mediated increase in sodium currents is normalized by Spike protein or NRP-1 inhibition in DRG neurons. Representative sodium current traces (**A, F**) recorded from small-sized DRGs neurons, incubated for 30 min with the indicated treatments, in response to depolarization steps from –70 to +60 mV from a holding potential of –60 mV. Summary of current-voltage curves (**B, G**) and normalized peak (**C, H**) currents (pA/pF) from DRG neurons as indicated. Boltzmann fits for normalized conductance G/G_max_ voltage relationship for voltage dependent activation (**D, I**) and inactivation (**E, J**) of the sensory neurons as indicated. Error bars indicate mean ± s.e.m. Half-maximal activation and inactivation (V_1/2_) and slope values (*k*) for activation and inactivation were not different between any of the conditions (p >0.9999, Kruskal-Wallis test with Dunn’s post hoc); values presented in Table S2. P values of comparisons between treatments are as indicated; for full statistical analyses see Table S1.

As calcium channels play multiple critical roles in the transmission and processing of pain-related information within the primary afferent pain pathway [61], we evaluated if they were affected. We focused on N-type (CaV2.2) channels as these mediate neurotransmitter release at afferent fiber synapses in the dorsal horn and are critical in the pain matrix [50]. VEGF-A facilitated a 1.8–fold increase in total Ca^2+^ currents compared to vehicle (PBS)-treated DRGs, which was completely blocked by Spike protein (Fig. 4A-C and Table S1). Spike protein alone did not affect Ca^2+^ currents (Fig. 4A-C). Additionally, we did not observe any changes in activation and inactivation kinetics between the conditions tested (Fig. 4D, E and Tables S1, S2). Similar results were obtained for the NRP-1 inhibitor EG00229, which inhibited the VEGF-A mediated increase in N-type Ca^2+^ currents (Fig. 4F-H and Table S1) but had no effect on the biophysical properties (Fig. 4I, J and Tables S1, S2). These data implicate Spike protein and NRP-1 in Na^+^ and Ca^2+^ (CaV2.2) channels in VEGF-A/NRP-1 signaling.

**Figure 4.**
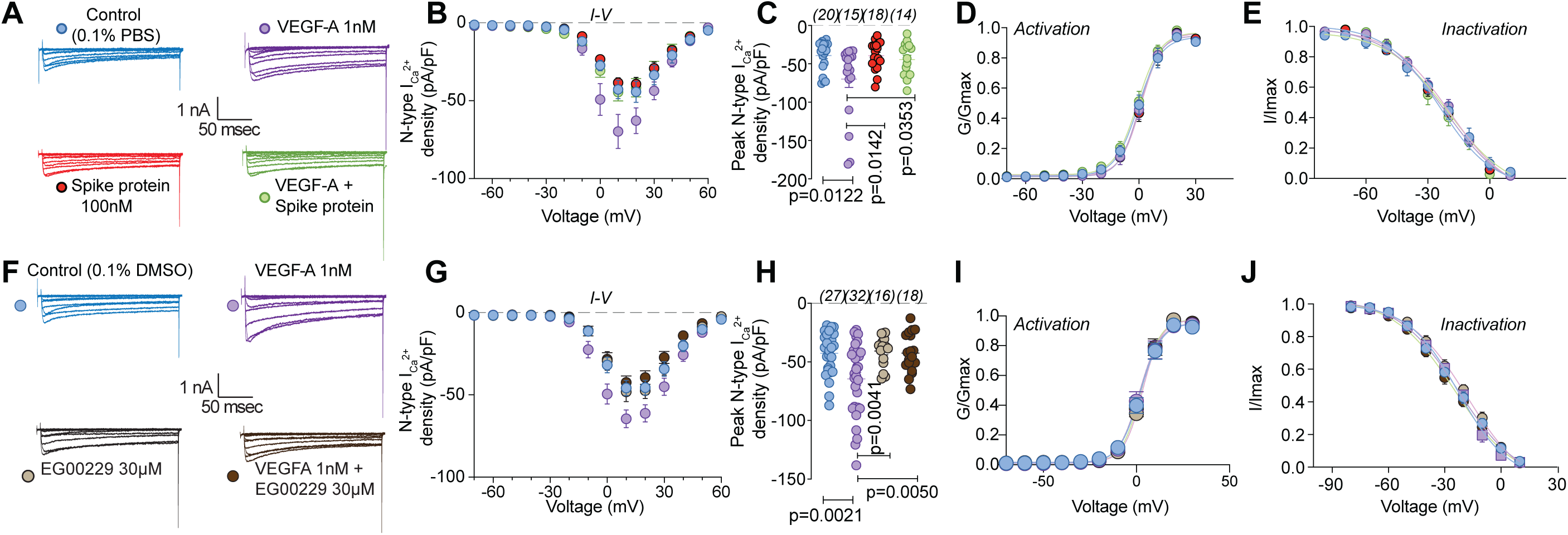
VEGF-A–mediated increase in calcium currents is normalized by Spike protein or NRP-1 inhibition in DRG neurons. Representative calcium current (via N-type channels) traces (**A, F**) recorded from small-sized DRGs neurons, incubated for 30 min with the indicated treatments, in response to holding voltage of –60 mV with 200-ms voltage steps applied at 5-s intervals in +10 mV increments from –70 to +60 mV. Pharmacological isolation of N-type (CaV2.2) current was achieved with a cocktail of toxins/small molecules. Summary of current-voltage curves (**B, G**) and normalized peak (**C, H**) currents (pA/pF) from DRG neurons as indicated. Boltzmann fits for normalized conductance G/Gmax voltage relations for voltage dependent activation (**D, I**) and inactivation (**E, J**) of the sensory neurons as indicated. Error bars indicate mean ± s.e.m. Half-maximal activation and inactivation (V_1/2_) and slope values (*k*) for activation and inactivation were not different between any of the conditions (p >0.9999, Kruskal-Wallis test with Dunn’s post hoc); values presented in Table S2. P values of comparisons between treatments are as indicated; for full statistical analyses see Table S1.

### 3.3. VEGF-A enhances synaptic activity in the lumbar dorsal horn that is normalized by inhibition of NRP-1 signaling and Spike protein

The spinal cord is an integrator of sensory transmission where incoming nociceptive signals undergo convergence and modulation [57]. Spinal presynaptic neurotransmission relies on DRG neuron action potential firing and neurotransmitter release. From these fundamental physiological principles, as well as the results described above, we were prompted to evaluate whether synaptic activity was affected in the lumbar dorsal horn. The amplitudes of spontaneous excitatory postsynaptic currents (sEPSCs) of neurons in the *substantia gelatinosa* region of the lumbar dorsal horn were not affected by VEGF-A (Fig. 5A, B and Table S1). In contrast, VEGF-A application increased sEPSC frequency by ∼3.6–fold, which was reduced by ∼57% by inhibition of NRP-1 with EG00229 and ∼50% by Spike protein (Fig. 5A, C and Table S1). Amplitude and inter-event interval cumulative distribution curves for sEPSCs are shown in Figure 5D, E. When compared to vehicle controls, VEGF-A, with or without NRP-1 inhibitor or Spike protein, had no effect on the cumulative amplitude distribution of the spontaneous EPSCs (Fig. 5D and Table S1) but changed the cumulative frequency distribution of spontaneous EPSCs with significantly longer inter-event intervals (Fig. 5E and Table S1). Together, these data suggest a presynaptic mechanism of action of Spike protein and NRP-1.

**Figure 5.**
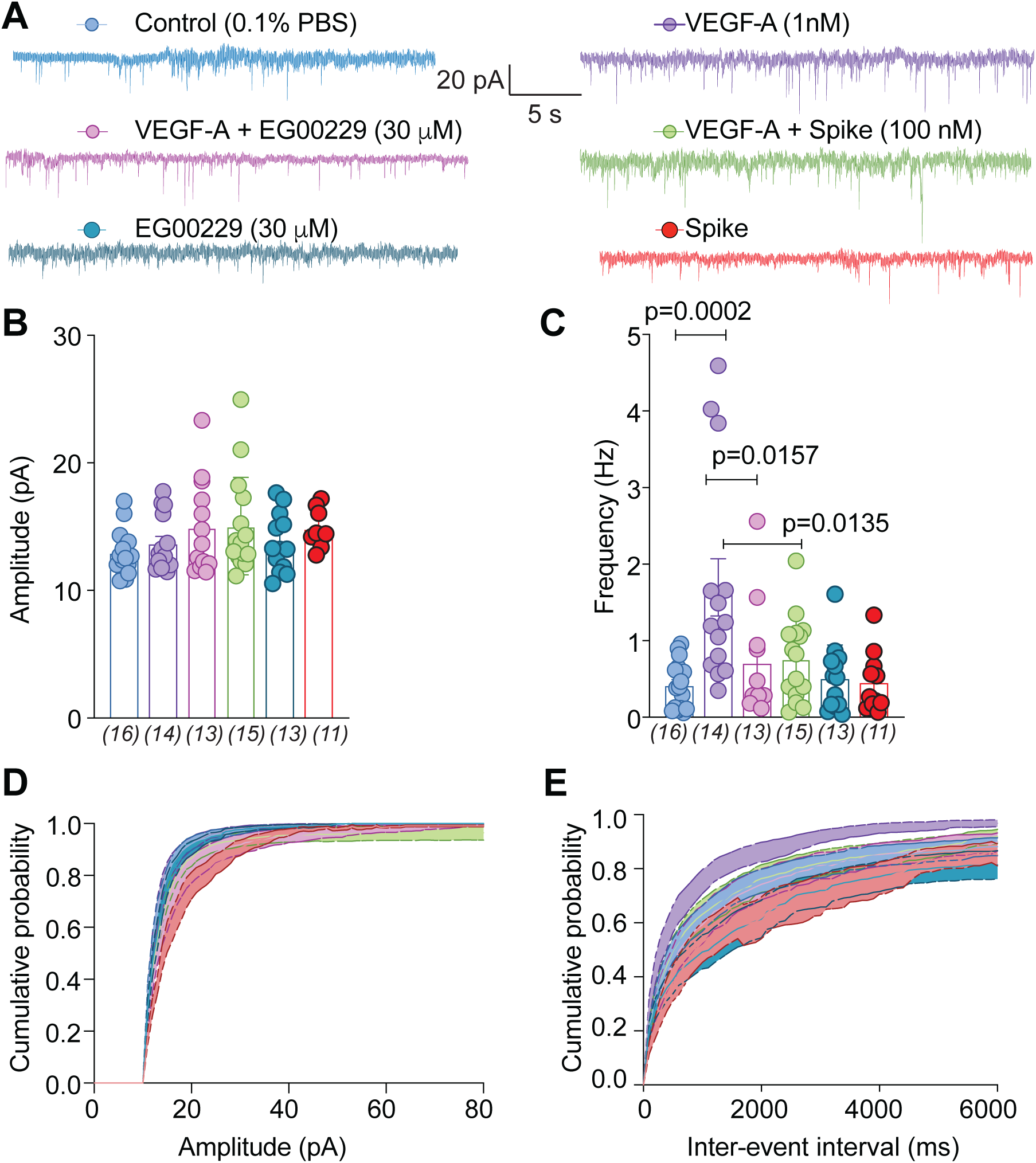
sEPSC frequency is reduced by pharmacological antagonism of NRP-1 by EG00229. (**A**) Representative traces of spontaneous excitatory postsynaptic currents (sEPSC) from neurons from the substantia gelatinosa in the superficial dorsal horn (lamina I/II) treated for at least 30 min with the indicated conditions. Summary of amplitudes (**B**) and frequencies (**C**) of sEPSCs for all groups are shown. Cumulative distribution of the sEPSCs amplitude (**D**) and the inter-event interval (**E**) recorded from cells as indicated. Perfusion of 30 μM EG00229 decreased spontaneous excitatory synaptic transmission (**A-E**) in lumbar dorsal horn neurons. P values of comparisons between treatments are as indicated; for full statistical analyses see Table S1.

### 3.5. Spike protein and inhibition of NRP-1 confer anti-nociception in the spared nerve injury model (SNI) of chronic neuropathic pain

We used the spared nerve injury (SNI) model of neuropathic pain, chosen because it produces a reliable and consistent increase in pain sensitivity [12], to evaluate the potential of disruption of the VEGF-A/NRP-1 pathway to reverse nociception. VEGF-A triggers autophosphorylation of VEGFR2 at Y1175 [54], thereby serving as a proxy for activation of VEGF-A signaling. In rats with SNI, intrathecal application of Spike, decreased the phosphorylation of VEGFR2 (Y1175) on both the contralateral (non-injured) and ipsilateral (injured) side (Fig. 6A, B). This shows that Spike can inhibit VEGF-A signaling in a rat model of chronic neuropathic pain. SNI injury efficiently reduced paw withdrawal thresholds (PWTs) (mechanical allodynia, Fig. 6C and Table S1) 10 days post injury. Spinal administration of Spike protein significantly increased PWTs (Fig. 6C and Table S1), in a dose-dependent manner, for 5 hours. Analysis of the area under the curve (AUC) confirmed the dose-dependent reversal of mechanical allodynia (Fig. 6D and Table S1) compared to vehicle-treated injured animals. Similar results were seen with female rats injected with Spike (Fig. 6E, F). Finally, inhibition of NRP-1 signaling with EG00229 also reversed paw-withdrawal thresholds (Fig. 6G, H and Table S1).

**Figure 6.**
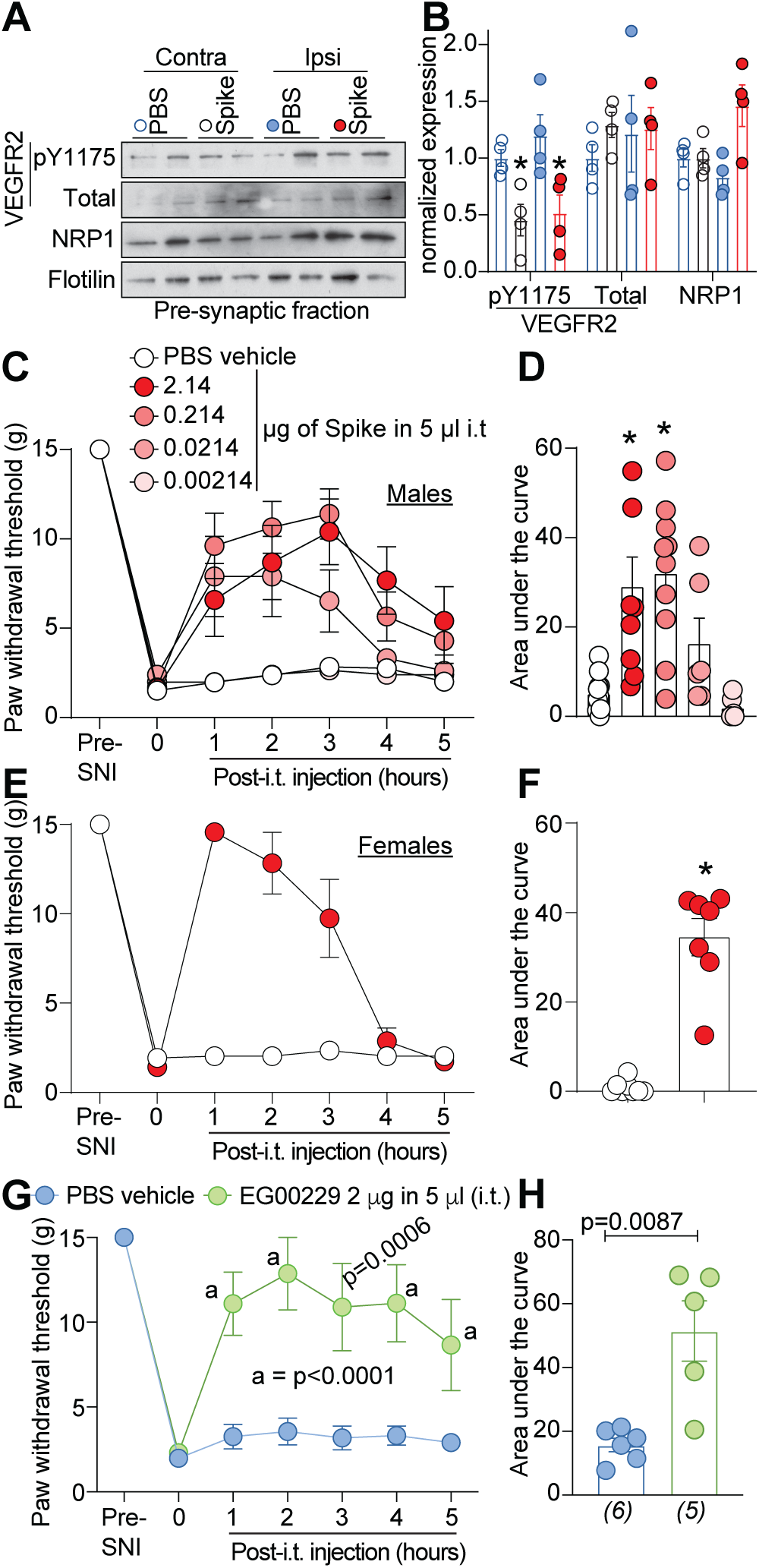
SARS-CoV-2’s Spike protein and NRP-1 antagonism reverses SNI-induced mechanical allodynia. Spared nerve injury (SNI) elicited mechanical allodynia 10 days after surgery. (**A**) Representative immunoblots of NRP-1, total and pY1175 VEGF-R2 levels at pre-synaptic sites in rat spinal dorsal horn after SNI. Tissues were collected 3 hours (i.e., at peak of anti-allodynia) following intrathecal injection of the receptor binding domain of the Spike protein (2.14 μg/5μl). (**B**) Bar graph with scatter plots showing the quantification of n= 4 samples as in **A** (*p<0.05, Kruskal-Wallis test). Paw withdrawal thresholds for SNI rats (male) administered saline (vehicle) or four doses of the receptor binding domain of the Spike protein (0.00214 to 2.14 μg/5μl) intrathecally (i.t.); n = 6-12) (**C**), 2.14 μg/5μl dose of Spike in female rats, or the NRP-1 inhibitor EG00229 in male rats (2 μg/5μl; n = 6) (**E**). (**D, F, H**) Summary of data shown in panels **C, E**, and **G** plotted as area under the curve (AUC) for 0-5 hours. P values, versus control, are indicated. Data is shown as mean ± s.e.m. and was analyzed by non-parametric two-way analysis of variance where time was the within subject factor and treatment was the between subject factor (post hoc: Sidak). AUCs were compared by Mann-Whitney test. The experiments were analyzed by an investigator blinded to the treatment. P values of comparisons between treatments are as indicated; for full statistical analyses see Table S1.

## 4.0. Discussion

Our data show that SARS-CoV-2 Spike protein subverts VEGF-A/NRP-1 pro-nociceptive signaling. Relevant to chronic neuropathic pain, Spike negated VEGF-A–mediated increases in: (*i*) voltage-gated sodium and N-type calcium current densities in DRG neurons; (*ii*) spontaneous firing in DRG neurons; (*iii*) spinal neurotransmission; and (*iv*) mechanical allodynia and thermal hyperalgesia. Consequently, Spike protein was analgesic in a nerve injury rat model. Based on the reported increase in VEGF-A levels in COVID-19 patients [30], one would expect to observe increased pain-related symptoms. However, our data suggest that the SARS-CoV-2 Spike protein hijacks NRP-1 signaling to ameliorate VEGF-A mediated pain. This raises the possibility that pain, as an early symptom of COVID-19, may be directly dampened by the SARS-CoV-2 Spike protein. Our results do not exclude the possibility that alternative fragments of Spike or other viral proteins may be pro-nociceptive. Leveraging this atypical function of SARS-CoV-2 Spike protein may yield a novel class of therapeutics for pain.

Clinical findings that VEGF-A contributes to pain are supported by observations that in osteoarthritis increased VEGF expression in synovial fluids has been associated with higher pain scores [55]. VEGF-A has been reported to enhance pain behaviors in normal, nerve-injured and diabetic animals [23; 58]. Blocking VEGF-A has been shown to reduce nociception in rodents and to exert a neuroprotective effect by improving neuronal restoration and conduction, decreasing pro-apoptotic Caspase-3 levels in sensory neurons, preventing neural perfusion and epidermal sensory fiber loss [24; 52; 67]. Alternative splicing of the VEGF-A gene produces several isoforms of the mature protein containing between 121 and 206 amino acid residues, with VEGF-A165a being pro-nociceptive [23] via sensitization of transient receptor potential (TRP) channels [24] and ATP-gated purinergic P2×2/3 receptors [27] in DRG neurons. VEGF-A165b is a VEGFR2 partial agonist that competes with VEGF-A165a for binding to VEGFR2. The VEGF-A165a isoform also binds to the NRP-1 co-receptor, whereas VEGFA165b does not [44]. Thus, it is a balance of the splice isoforms (VEGF-Aa/b) that likely determines the nociceptive impact effect on sensory neurons.

Our data shows that VEGF-A elicits long-lasting (up to 24 hours) mechanical allodynia and thermal hyperalgesia in male and female naïve rats, thus supporting the premise that VEGF-A is pro-nociceptive. VEGF is augmented in serum of rheumatoid arthritis patients [4; 40]. The levels of VEGF-A were substantially lower in the sera of asymptomatic individuals compared to symptomatic individuals and matched those found in healthy controls [30]. Conversely, transcript levels of the VEGF-A co-receptor NRP-1 were increased in COVID-19 patients compared to healthy controls [6]. NRP-1 is a dimeric transmembrane receptor that regulates pleiotropic biological processes, including axon guidance, angiogenesis and vascular permeability [21; 45; 56]. In chronic neuropathic pain, a concomitant increase of NRP-1 and VEGF-A have been reported in DRG neurons [29]. We observed that pharmacological antagonism of the VEGF-A binding b1b2 domain of NRP-1 using EG00229 resulted in decreased mechanical allodynia in SNI. A corollary to this, VEGF-A–induced DRG sensitization and the related allodynia/hyperalgesia in naïve rats was annulled by NRP-1 antagonism. This work identifies a heretofore unknown role of VEGF-A/NRP-1 signaling in pain. Our working model shows that VEGF-A engagement of NRP-1 is blocked by Spike protein consequently decreasing activities of two key nociceptive voltage-gated sodium, likely NaV1.7, and calcium (CaV2.2) channels (Fig. 1D). The resulting decrease in spontaneous DRG neuronal firing by Spike protein translates into a reduction in pain (Fig. 1D). Characterization of the molecular cascade downstream of VEGF-A/NRP-1 signaling awaits further work.

Altogether, our data suggest that interfering with VEGF-A/NRP-1 using SARS-CoV-2 Spike or the NRP-1 inhibitor EG00229 is analgesic. For cancer, this pathway has been extensively targeted for anti-angiogenesis. A monoclonal antibody targeting VEGF-A (bevacizumab, Avastin®) has been used as a cancer treatment [20]. A phase 1a clinical trial of NRP-1 antibody MNRP1685A (Vesencumab®), targeting the against VEGF-binding site of NRP-1, was well-tolerated in cancer patients [62], but neuropathy was noted. Possible reasons for this include the possibility that anti-VEGF-A therapies may target the alternatively spliced VEGF-A165b isoform, which has a neuroprotective role. It is important to note that the VEGF-A165b isoform binds to the VEGFR2 binding site with equal affinity to VEGF165a but does not bind to the co-receptor neuropilin 1 (NRP-1) [7], thus acting as an inhibitor of the VEGF-A-NRP-1 pathway [3]. Anti-VEGF-A therapies may simultaneously inhibit both pro-and anti-algesic pathways involving the multiple VEGF-A isoforms, thus an ensuing loss of balance may favor nociceptor sensitization. Here, we are specifically studying the downstream actions following the binding of VEGF-A165a to the b1 domain of NRP-1. Targeting this interaction may be more specific compared to broad VEGF-A inhibitors that have been developed for cancer treatment. Thus, our preclinical work provides a rationale for targeting the VEGF-A/NRP-1 pro-nociceptive signaling axis in future clinical trials.

## ABBREVIATIONS USED

ACE2: Angiotensin converting enzyme 2
AUC: area under the curve
CaV2.2: N-type voltage-gated calcium channel
COVID-19: coronavirus disease 2019
DRG: dorsal root ganglia
MEA: multi-well microelectrode array
NaV1.7: voltage-gated sodium channel isoform 7
NRP-1: Neuropilin-1
PWTs: paw withdrawal thresholds
SARS-CoV-2: Severe acute respiratory syndrome coronavirus 2
sEPSCs: spontaneous excitatory postsynaptic currents
SNI: spared nerve injury
VEGF-A: vascular endothelial growth factor-A

## ACKNOWLEDGMENTS

We thank Dr. Marcel Patek (BrightRock Path LLC) for critically reading the manuscript.

## Author contributions

R.K. and A.M. developed the concept and designed experiments; A.M., L.F.M., L.B., K.G., D.R., Y.Z., H.J.S., S.C., S.L., K.B.G., and S.P.-M. collected and analyzed data; L.F.M., K.B.G., and S.L. performed animal behavior studies; L.B., K.G., D.R., Y.Z., H.J.S., and S.C. performed electrophysiology recordings; S.P.-M. assisted with docking studies; A.P. and M.M.I. provided funding for L.F.M.; R.K. and A.M. wrote the manuscript; and R.K. and A.M. supervised all aspects of this project. All authors had the opportunity to discuss results and comment on the manuscript;

## Data and materials availability

All data is available in the main text, figures, and supplementary materials.

