## Supplementary Materials for "SARS-CoV-2 Spike protein co-opts VEGF-A/Neuropilin-1 receptor signaling to induce analgesia"

^†^Contributed equally

**This PDF file includes:**

Materials and Methods

Tables S1 to S2

References (41-51)

Materials and Methods

Animals: Pathogen-free, adult male (250g) or female (225g) Sprague–Dawley rats (Envigo) were housed in temperature (23 ± 3 °C) and light (12-h light/12-h dark cycle; lights on 07:00–19:00) controlled rooms with standard rodent chow and water available ad libitum. The Institutional Animal Care and Use Committee of the College of Medicine at the University of Arizona approved all experiments. All procedures were conducted in accordance with the Guide for Care and Use of Laboratory Animals published by the National Institutes of Health and the ethical guidelines of the International Association for the Study of Pain. Animals were randomly assigned to treatment or control groups for the behavioral studies. Animals were initially housed three per cage but individually housed after the intrathecal cannulation on a 12 h light-dark cycle with food and water ad libitum. All behavioral experiments were performed by experimenters who were blinded to the experimental groups and treatments.

Preparation of cultured dorsal root ganglia neurons: Female Sprague–Dawley rats (100 g; Envigo) were deeply anaesthetized with isoflurane overdose (5% in air) and sacrificed by rapid decapitation. Following laminectomy, dorsal root ganglia (DRG) were quickly removed, trimmed at their roots, and enzymatically digested in 3 mL bicarbonate-free, serum-free, sterile DMEM (Cat# 11965, Thermo Fisher Scientific, Waltham, MA) solution containing neutral protease (3.125 mg/mL, Cat#LS02104; Worthington, Lakewood, NJ) and collagenase type I (5 mg/mL, Cat# LS004194, Worthington, Lakewood, NJ). Subsequently, the isolated DRGs were incubated with the enzyme cocktail for 60 minutes at 37˚C under gentle agitation. The digested DRGs were then mechanically separated by gently passing them through the tip of a 1 mL pipette until a single cell suspension was obtained. The fully dissociated DRG neurons were then gently centrifuged to collect the cells (~1.5 x 10^6^) as a pellet and the supernatant was discarded. The cells were resuspended and washed with DRG media (DMEM containing 1% penicillin/streptomycin sulfate from 10,000 μg/mL stock, and 10% fetal bovine serum (Hyclone)) before plating onto poly-D-lysine and laminin-coated 12-mm glass coverslips. All whole-cell electrophysiology experiments were performed within 48 h of plating DRG neurons since electrophysiological profiles change during this period.

Whole-cell electrophysiological recordings of sodium and calcium currents in cultured rat DRG neurons: All recordings were obtained from acutely dissociated DRG neurons from Sprague Dawley rats, using procedures adapted from our prior work [64; 67; 69]. For sodium current recordings the internal pipette solution consisted of (in mM): 140 CsF, 10 NaCl, 1.1Cs-EGTA, and 15 HEPES (pH 7.3, mOsm/L = 290-310) and external solution contained (in mM): 140 NaCl, 30 tetraethylammonium chloride, 10 D-glucose, 3 KCl, 1 CaCl_2_, 0.5 CdCl_2_, 1 MgCl_2_, and 10 HEPES (pH 7.3, mOsm/L = 310-315). DRG neurons were interrogated with current-voltage (I-V) and activation/inactivation voltage protocols as described previously [63; 65]. The voltage protocols were as follows: (a) I-V protocol: from a holding potential of −60 mV, cells were depolarized with 150-millisecond voltage steps over a range of −70 to +60 mV in +5-mV increments. This permitted acquisition of current density values such that the activation of sodium channels, occurring between ~0 to 10 mV, could be analyzed as a function of voltage, from which peak current density was inferred (normalized to cell capacitance (in picofarads, pF)); (b) inactivation protocol: from a holding potential of −60 mV, cells were subjected to hyperpolarizing/repolarizing pulses for 1 second over a range of −120 to 0 mV in +10 mV steps. This incremental increase in membrane potential conditioned various proportions of sodium channels into a state of fast-inactivation – in this case the 0-mV test pulse for 200 milliseconds revealed fast inactivation when normalized to maximum sodium current [63].

Recordings of N-type (CaV2.2) voltage-gated calcium currents were obtained using recording solutions and protocols described earlier [71]. The intracellular pipette solution was composed of (in mM): 150 CsCl_2_, 10 HEPES, 5 Mg-ATP, and 5 BAPTA (pH 7.3, mOsm/L=290-310) and the external solution contained (in mM): 110 NMDG, 10 BaCl_2_, 30 TEA-Cl, 10 HEPES, 10 glucose and 1 μM TTX (pH 7.3, mOsm/L = ~ 310). To isolate N-type specific calcium currents, the following blockers were used: SNX482 (200 nM, R-type Ca^2+^ channel blocker), TTA-P2 (1 μM, T-type Ca^2+^ channel blocker), ω-agatoxin (200 nM, P/Q-type Ca^2+^ channel blocker), and nifedipine (10 μM, L-type Ca^2+^ channel blocker). Activation of I_Ca_ was measured from a holding voltage of -60 mV for 5 ms followed by 200-ms depolarizing voltage steps from -70 mV to +60 mV in 10-mV increments. Whole-cell currents were normalized to cellular capacitance for analysis of channel activation profiles as a function of voltage in addition to peak current density. Steady-state inactivation of I_Ca_ was determined by applying a 1500 ms conditioning prepulse (−100 to +30 mV in +10 mV increments) after which, the voltage was stepped to +10 mV for 200-ms. There were 15-s intervals separating each acquisition to allow channels to revert to their basal state.

Pipettes with 1 to 3 MΩ resistance were used for all recordings and pulled from borosilicate capillaries on a Flaming/Brown P-97 puller (Sutter Instruments, California).

Hind paw injection procedures: PBS vehicle (NaCl 137 mM, KCl 2.5 mM, Na_2_HPO_4_ 10 mM and KH_2_PO_4_ 1.8 mM), VEGF-A_165_ (10 nM), Spike (1 µM) and EG00229 [68] (30 µM, Cat. No. 6986, Tocris Bioscience) were injected subcutaneously, alone or in combination, in the dorsum of the left hind paw. Rats were gently restrained under a fabric cloth, and 50 µL were injected using 0.5 mL syringes (27-G needles).

Preparation of Spinal Cord slices: Pathogen-free, male Sprague-Dawley rat pups (10-15 days old; Envigo) were deeply anesthetized with isoflurane (4% for induction and 2% for maintaining). For spinal nerve block, 0.3 mL of 2% lidocaine was injected to both sides of L4 to 5 lumbar vertebrae. Laminectomy was performed from mid-thoracic to low lumbar levels, and the spinal cord was quickly removed to cold modified ACSF oxygenated with 95% O2 and 5% CO2. The ACSF for dissection contained the following (in millimolar): 80 NaCl, 2.5 KCl, 1.25 NaH2PO4, 0.5 CaCl2.2H_2_O, 3.5 MgCl2.6H_2_O, 25 NaHCO3, 75 sucrose, 1.3 ascorbate, 3.0 sodium pyruvate, with pH at 7.4 and osmolarity at 310 mOsm. Transverse 380-mm thick slices were obtained by a vibratome (VT1200S; Leica, Nussloch, Germany). Slices were then incubated for at least 40 min at 37°C and then for 1h at RT in an oxygenated recording solution containing the following (in millimolar): 125 NaCl, 2.5 KCl, 2 CaCl2.2H_2_O, 1 MgCl2.6H_2_O, 1.25 NaH2PO4, 26 NaHCO3, 25 D-glucose, 1.3 ascorbate, 3.0 sodium pyruvate, with pH at 7.4 and osmolarity at 320 mOsm. The slices were then positioned in a recording chamber and continuously perfused with oxygenated recording solution at a rate of 3 to 4 mL/min before electrophysiological recordings at RT.

Electrophysiological recordings in spinal cord slices by whole-cell patch clamp: *Substantia gelatinosa* neurons (lamina I/II) were visualized and identified in the slices by means of infrared differential interference contrast video microscopy on an upright microscope (FN1; Nikon, Tokyo, Japan) equipped with a 3.40/0.80 water-immersion objective and a charge-coupled device camera. Patch pipettes with resistance at 6 to 10MΩ were made from borosilicate glass (Sutter Instruments, Novato, CA) on a four-step micropipette puller (P-90; Sutter Instruments, Novato, CA). The pipette solution contained the following (in millimolar): 120 potassium gluconate, 20 KCl, 2 MgCl2, 2Na2-ATP, 0.5 Na-GTP, 20 HEPES, 0.5 EGTA, with pH at 7.28 and osmolarity at 310 mOsm. The membrane potential was held at − 60 mV using a PATCHMASTER software in combination with a patch clamp amplifier (EPC10; HEKA Elektronik, Lambrecht, Germany). The whole-cell configuration was obtained in voltage-clamp mode. To record spontaneous excitatory postsynaptic currents (sEPSCs), bicuculline methiodide (10 μM) and strychnine (2 μM) were added to the recording solution to block γ-aminobutyric acid-activated (GABA) and glycine-activated currents. VEGFA (1nM), NRP-1 inhibitor (EG00229, 30 µM) and Spike protein were added directly to the recording solution as indicated.

Hyperpolarizing step pulses (5 mV in intensity, 50 milliseconds in duration) were periodically delivered to monitor the access resistance (15–25 MΩ), and recordings were discontinued if the access resistance changed by more than 20%. For each neuron, sEPSCs were recorded for a total duration of 2 min. Currents were filtered at 3 kHz and digitized at 5 kHz. Data were further analyzed by the Mini-Analysis Program (Synatosoft Inc., NJ) to provide spreadsheets for the generation of cumulative probability plots. The amplitude and frequency of sEPSCs were compared between neurons from animals in control and the indicated groups.

Synapse enrichment and fractionation: Adult rats were killed by isofluorane overdose and decapitation, the spinal cords dissected, the lumbar region isolated and separated into contralateral and ipsilateral sides. Only the dorsal horn of the spinal cord was used as this structure contains the synapses arising from the DRG. Synaptosomes isolation was done according to [70]. Fresh tissues were homogenized in ice-cold Sucrose 0.32M, HEPES 10 mM, pH 7.4 buffer. The homogenates were centrifuged at 1000xg for 10 min at 4°C to pellet the insoluble material. The supernatant was harvested and centrifuged at 12000xg for 20 min at 4°C to pellet a crude membrane fraction. The pellet was then re-suspended in a hypotonic buffer (4 mM HEPES, 1 mM EDTA, pH 7.4) and the resulting synaptosomes pelleted by centrifugation at 12000xg for 20 min at 4°C. The synaptosomes were then incubated in 20 mM HEPES, 100 mM NaCl, 0.5% triton X, pH= 7.2) for 15 min on ice and centrifuged at 12000xg for 20 min at 4°C. The supernatant was considered as the non-postsynaptic density (non-PSD) membrane fraction, sometimes referred to as the triton soluble fraction. All buffers were supplemented with protease (Cat#B14002) and phosphatase (Cat#B15002) inhibitor cocktails (Bimake).

Implantation of intrathecal catheter: For intrathecal drug administration, rats were chronically implanted with catheters as described (Yaksh and Rudy, 1976). Rats were anesthetized (ketamine/xylazine anesthesia, 80/12 mg/kg i.p ) and placed in a stereotaxic head holder, the occipital muscles were separated from their occipital insertion and retracted caudally to expose the cisternal membrane at the base of the skull, the cisterna magna was exposed and incised, an 8-cm catheter (PE10 polyethylene tubing) was passed caudally from the cisterna magna to the level of the lumbar enlargement. Catheters were sutured (3-0 silk suture) into the deep muscle and externalized at the back of the neck; skin was closed with auto clips. Animals were allowed to recover and were examined for evidence of neurologic injury. Animals with evidence of neuromuscular deficits were excluded.

Spared nerve injury (SNI): After a recovery period of 7 days after implantation of intrathecal catheter, the spared nerve injury was induced. Under isoflurane anesthesia (5% induction, 2.0% maintenance in 2 L/min air), skin on the lateral surface of the left hind thigh was incised. The biceps femoris muscle was bluntly dissected to expose the three terminal branches of the sciatic nerve [62]. Briefly, the common peroneal and tibial branches were tightly ligated with 5-0 silk, 2–3 mm of the nerves was removed below the ligations, with special care taken to avoid any damage to the sural nerve. Closure of the incision was made in two layers. The muscle was sutured once with 3-0 silk suture; skin was auto-clipped. Animals were allowed to recover for 12-14 days before the drug testing.

Tactile sensory thresholds: The assessment of tactile allodynia (i.e., a decreased threshold for paw withdrawal after probing with normally innocuous mechanical stimuli) consisted of testing the withdrawal threshold of the paw in response to probing with a series of calibrated fine (von Frey) filaments. Each filament was applied perpendicularly to the plantar surface of the paw of rats held in suspended wire mesh cages. We determined the withdrawal threshold by sequentially increasing and decreasing the stimulus strength (the ‘up and down’ method), and we analyzed data using the nonparametric method of Dixon (as described by Chaplan et al. ) with results expressed as the mean withdrawal threshold

Thermal sensory thresholds: Paw withdrawal latencies were determined as described by Hargreaves et al.[66] was used. Rats were acclimated within Plexiglas enclosures on a clear glass plate for 30 minutes. A radiant heat source (high-intensity projector lamp) was focused onto the plantar surface of the hind paw. A motion detector halted the stimulus and a timer when the paw was withdrawn. To prevent tissue damage, a maximal cutoff of 33.5 sec was used.

Statistical analyses: All data was first tested for a Gaussian distribution using a D’Agostino-Pearson test (Prism 8 Software, Graphpad, San Diego, CA). The statistical significance of differences between means was determined by a parametric ANOVA followed by Tukey’s post hoc or a non-parametric Kruskal Wallis test followed by Dunn’s post-hoc test depending on if datasets achieved normality. Behavioral data with a time course were analyzed by Two-way ANOVA with Sidak’s post hoc test. Differences were considered significant if p≤ 0.05. Error bars in the graphs represent mean ± SEM. Full statistical analyses are described in Table 1. All data were plotted in Prism 8.

**Table S1. Statistical analyses of experiments.**

| **Figure panel** | **Assay** | **Statistical test; findings** | **Post-hoc analysis**  **(adjusted p-values)** | **Number of subjects** | **Number of subjects excluded (ROUT test)** |
| --- | --- | --- | --- | --- | --- |
| Figure 1A | multiwell microelectrode arrays (MEAs) on dorsal root ganglion sensory neurons – mean firing rate (Hz) | One-way ANOVA  p <0.0001 | Holm- Sidak's multiple comparisons test  PBS vs. Sema 3A p = 0.5439;  PBS vs. VEGF-A165  p<0.0001;  PBS vs. VEGF-B p = 0.5439;  VEGF-A165 vs. Spike protein 100nM p<0.0001;  VEGF-A165 vs. EG p<0.0001 | PBS n = 56;  Sema 3A n = 42;  VEGF-B n = 37;  VEGF-A n = 42;  VEGF-A + Spike protein n = 95;  VEGF-A + EG00229 n = 90 | None |
| Figure 2A | Naïve male rats – paw withdrawal threshold | Two-way ANOVA  p <0.0001 | Sidak's multiple comparisons test  Time after injection:  0.5 h  PBS vs. VEGF-A165 p<0.0001  PBS vs. Spike p>0.9999  PBS vs. EG00229 p>0.9999  PBS vs. EG00229 + VEGF-A165 p=0.9581  PBS vs. VEGF-A165 + Spike p>0.9999  1 h  PBS vs. VEGF-A165 p<0.0001  PBS vs. Spike p=0.997  PBS vs. EG00229 p=0.9997  PBS vs. EG00229 + VEGF-A165 p=0.0273  PBS vs. VEGF-A165 + Spike p=0.9891  2 h  PBS vs. VEGF-A165 p=<0.0001  PBS vs. Spike p>0.9999  PBS vs. EG00229 p>0.9999  PBS vs. EG00229 + VEGF-A165 p<0.0001  PBS vs. VEGF-A165 + Spike p>0.9999  3 h  PBS vs. VEGF-A165 p<0.0001  PBS vs. Spike p>0.9999  PBS vs. EG00229 p>0.9999  PBS vs. EG00229 + VEGF-A165 p<0.0001  PBS vs. VEGF-A165 + Spike p=0.0586  4 h  PBS vs. VEGF-A165 p<0.0001  PBS vs. Spike p>0.9999  PBS vs. EG00229 p>0.9999  PBS vs. EG00229 + VEGF-A165 p=0.0002  PBS vs. VEGF-A165 + Spike p=0.0066  5 h  PBS vs. VEGF-A165 p<0.0001  PBS vs. Spike p>0.9999  PBS vs. EG00229 p>0.9999  PBS vs. EG00229 + VEGF-A165 p<0.0001  PBS vs. VEGF-A165 + Spike p=0.0001  6 h  PBS vs. VEGF-A165 p<0.0001  PBS vs. Spike p>0.9999  PBS vs. EG00229 p>0.9999  PBS vs. EG00229 + VEGF-A165 p <0.0001  PBS vs. VEGF-A165 + Spike p= 0.0009    7 h  PBS vs. VEGF-A165 p<0.0001  PBS vs. Spike p>0.9999  PBS vs. EG00229 p>0.9999  PBS vs. EG00229 + VEGF-A165 p= 0.0029  PBS vs. VEGF-A165 + Spike p <0.0001    9 h  PBS vs. VEGF-A165 p<0.0001  PBS vs. Spike p>0.9999  PBS vs. EG00229 p>0.9999  PBS vs. EG00229 + VEGF-A165 p= 0.0018  PBS vs. VEGF-A165 + Spike p= 0.0003 | PBS n = 12;  VEGF-A n = 12;  Spike protein n = 6;  EG00229 n = 6;  VEGF-A + EG00229 n = 6;  VEGF-A + Spike protein n = 6; | None |
| Figure 2B | Naïve male rats – paw withdrawal threshold: Area over the curve | One-way ANOVA  p <0.0001 | Sidak's multiple comparisons test  PBS vs. VEGF-A165 p<0.0001  PBS vs. Spike p=0.9997  PBS vs. EG00229 p>0.9999  PBS vs. EG00229 + VEGFA165 p<0.0001  PBS vs. VEGFA165 + Spike p=0.0111  VEGF-A165 vs. EG00229 + VEGFA165 p=0.0004  VEGF-A165 vs. VEGFA165 + Spike p<0.0001 | PBS n = 12;  VEGF-A n = 12;  Spike protein n = 6;  EG00229 n = 6;  VEGF-A + EG00229 n = 6;  VEGF-A + Spike protein n = 6; | None |
| Figure 2C | Naïve male rats – paw withdrawal latency | Two-way ANOVA  p <0.0001 | Sidak's multiple comparisons test  Time after injection:  0.5 h  PBS vs. VEGF-A165 p=0.8005  PBS vs. Spike p>0.9999  PBS vs. EG00229 p>0.9999  PBS vs. EG00229 + VEGF-A165 p=0.9999  PBS vs. VEGFA165 + Spike p=0.9996    1 h  PBS vs. VEGF-A165 p<0.0001  PBS vs. Spike p>0.9999  PBS vs. EG00229 p>0.9999  PBS vs. EG00229 + VEGF-A165 p= 0.1568  PBS vs. VEGFA165 + Spike p>0.9999    2 h  PBS vs. VEGF-A165 p<0.0001  PBS vs. Spike p>0.9999  PBS vs. EG00229 p>0.9999  PBS vs. EG00229 + VEGF-A165 p= 0.0017  PBS vs. VEGFA165 + Spike p= 0.04    3 h  PBS vs. VEGF-A165 p<0.0001  PBS vs. Spike p=0.9993  PBS vs. EG00229 p>0.9999  PBS vs. EG00229 + VEGF-A165 p=0.0001  PBS vs. VEGFA165 + Spike p=0.0087    4 h  PBS vs. VEGF-A165 p<0.0001  PBS vs. Spike p>0.9999  PBS vs. EG00229 p>0.9999  PBS vs. EG00229 + VEGF-A165 p<0.0001  PBS vs. VEGFA165 + Spike p=0.3511    5 h  PBS vs. VEGF-A165 p<0.0001  PBS vs. Spike p>0.9999  PBS vs. EG00229 p>0.9999  PBS vs. EG00229 + VEGF-A165 p=0.0002  PBS vs. VEGFA165 + Spike p=0.1723    6 h  PBS vs. VEGF-A165 p<0.0001  PBS vs. Spike p>0.9999  PBS vs. EG00229 p>0.9999  PBS vs. EG00229 + VEGF-A165 p=0.0052  PBS vs. VEGFA165 + Spike p=0.4168    7 h  PBS vs. VEGF-A165 p<0.0001  PBS vs. Spike p>0.9999  PBS vs. EG00229 p= .9993  PBS vs. EG00229 + VEGF-A165 p<0.0001  PBS vs. VEGFA165 + Spike p=0.0774    9 h  PBS vs. VEGF-A165 p<0.0001  PBS vs. Spike p>0.9999  PBS vs. EG00229 p>0.9999  PBS vs. EG00229 + VEGF-A165 p=0.0012  PBS vs. VEGFA165 + Spike p=0.0202 | PBS n = 12;  VEGF-A n = 12;  Spike protein n = 6;  EG00229 n = 6;  VEGF-A + EG00229 n = 6;  VEGF-A + Spike protein n = 6; | None |
| Figure 2D | Naïve male rats – paw withdrawal latency: Area over the curve | One-way ANOVA  p <0.0001 | Sidak's multiple comparisons test  PBS vs. VEGF-A165 p<0.0001  PBS vs. Spike p=0.9776  PBS vs. EG00229 p>0.9999  PBS vs. EG00229 + VEGFA165 p<0.0001  PBS vs. VEGFA165 + Spike p=0.0039  VEGF-A165 vs. EG00229 + VEGFA165 p=0.4728  VEGF-A165 vs. VEGFA165 + Spike p<0.0001 | PBS n = 12;  VEGF-A n = 12;  Spike protein n = 6;  EG00229 n = 6;  VEGF-A + EG00229 n = 6;  VEGF-A + Spike protein n = 6; |  |
| Figure 2E | Naïve female rats – paw withdrawal threshold | Two-way ANOVA  p <0.0001 | Sidak's multiple comparisons test  Time after injection:  0.5 h  PBS vs. VEGF-A165 p<0.0001  PBS vs. Spike p>0.9999  PBS vs. EG00229 p>0.9999  PBS vs. EG00229 + VEGF-A165 p=0.9902  PBS vs. VEGF-A165 + Spike p>0.9999  1 h  PBS vs. VEGF-A165 p<0.0001  PBS vs. Spike p>0.9999  PBS vs. EG00229 p>0.9999  PBS vs. EG00229 + VEGF-A165 p=0.9709  PBS vs. VEGF-A165 + Spike p=0.1298  2 h  PBS vs. VEGF-A165 p=<0.0001  PBS vs. Spike p>0.9999  PBS vs. EG00229 p>0.9999  PBS vs. EG00229 + VEGF-A165 p=0.0038  PBS vs. VEGF-A165 + Spike p=0.2143  3 h  PBS vs. VEGF-A165 p<0.0001  PBS vs. Spike p>0.9999  PBS vs. EG00229 p>0.9999  PBS vs. EG00229 + VEGF-A165 p=0.1133  PBS vs. VEGF-A165 + Spike p>0.9999  4 h  PBS vs. VEGF-A165 p=0.0004  PBS vs. Spike p>0.9999  PBS vs. EG00229 p=0.9993  PBS vs. EG00229 + VEGF-A165 p=0.0012  PBS vs. VEGF-A165 + Spike p=0.6620  5 h  PBS vs. VEGF-A165 p<0.0001  PBS vs. Spike p>0.9999  PBS vs. EG00229 p>0.9999  PBS vs. EG00229 + VEGF-A165 p<0.0001  PBS vs. VEGF-A165 + Spike p=0.5423  6 h  PBS vs. VEGF-A165 p<0.0001  PBS vs. Spike p>0.9999  PBS vs. EG00229 p>0.9999  PBS vs. EG00229 + VEGF-A165 p<0.0001  PBS vs. VEGF-A165 + Spike p=0.1275    7 h  PBS vs. VEGF-A165 p=0.0113  PBS vs. Spike p>0.9999  PBS vs. EG00229 p>0.9999  PBS vs. EG00229 + VEGF-A165 p=0.0039  PBS vs. VEGF-A165 + Spike p>0.9999    9 h  PBS vs. VEGF-A165 p<0.0001  PBS vs. Spike p>0.9999  PBS vs. EG00229 p>0.9999  PBS vs. EG00229 + VEGF-A165 p=0.0311  PBS vs. VEGF-A165 + Spike p=0.1095 | PBS n = 12;  VEGF-A n = 12;  Spike protein n = 6;  EG00229 n = 6;  VEGF-A + EG00229 n = 6;  VEGF-A + Spike protein n = 6; | None |
| Figure 2F | Naïve female rats – paw withdrawal threshold: Area over the curve | One-way ANOVA  p <0.0001 | Sidak's multiple comparisons test  PBS vs. VEGF-A165 p<0.0001  PBS vs. Spike p=0.9516  PBS vs. EG00229 p=0.8341  PBS vs. EG00229 + VEGFA165 p=0.0066  PBS vs. VEGFA165 + Spike p=0.0810  VEGF-A165 vs. EG00229 + VEGFA165 p=0.2626  VEGF-A165 vs. VEGFA165 + Spike p=0.0320 | PBS n = 12;  VEGF-A n = 12;  Spike protein n = 6;  EG00229 n = 6;  VEGF-A + EG00229 n = 6;  VEGF-A + Spike protein n = 6; | None |
| Figure 2G | Naïve female rats – paw withdrawal latency | Two-way ANOVA  p <0.0001 | Sidak's multiple comparisons test  Time after injection:  0.5 h  PBS vs. VEGF-A165 p<0.0001  PBS vs. Spike p>0.9999  PBS vs. EG00229 p=0.9984  PBS vs. EG00229 + VEGF-A165 p=0.8339  PBS vs. VEGFA165 + Spike p=0.9984    1 h  PBS vs. VEGF-A165 p<0.0001  PBS vs. Spike p>0.9999  PBS vs. EG00229 p>0.9999  PBS vs. EG00229 + VEGF-A165 p>0.9999  PBS vs. VEGFA165 + Spike p=0.3044    2 h  PBS vs. VEGF-A165 p<0.0001  PBS vs. Spike p>0.9999  PBS vs. EG00229 p>0.9999  PBS vs. EG00229 + VEGF-A165 p=0.1905  PBS vs. VEGFA165 + Spike p=0.1617    3 h  PBS vs. VEGF-A165 p=0.0848  PBS vs. Spike p>0.9999  PBS vs. EG00229 p>0.9999  PBS vs. EG00229 + VEGF-A165 p=0.3859  PBS vs. VEGFA165 + Spike p=0.1404    4 h  PBS vs. VEGF-A165 p=0.3047  PBS vs. Spike p>0.9999  PBS vs. EG00229 p>0.9999  PBS vs. EG00229 + VEGF-A165 p=0.0471  PBS vs. VEGFA165 + Spike p=0.9958    5 h  PBS vs. VEGF-A165 p=0.1732  PBS vs. Spike p>0.9999  PBS vs. EG00229 p>0.9999  PBS vs. EG00229 + VEGF-A165 p=0.0349  PBS vs. VEGFA165 + Spike p=0.9126    6 h  PBS vs. VEGF-A165 p=0.0111  PBS vs. Spike p>0.9999  PBS vs. EG00229 p>0.9999  PBS vs. EG00229 + VEGF-A165 p=0.0412  PBS vs. VEGFA165 + Spike p=0.0943    7 h  PBS vs. VEGF-A165 p=0.0088  PBS vs. Spike p=0.9998  PBS vs. EG00229 p=0.9996  PBS vs. EG00229 + VEGF-A165 p=0.0761  PBS vs. VEGFA165 + Spike p=0.0536    9 h  PBS vs. VEGF-A165 p=0.0348  PBS vs. Spike p>0.9999  PBS vs. EG00229 p>0.9999  PBS vs. EG00229 + VEGF-A165 p=0.0011  PBS vs. VEGFA165 + Spike p=0.3535 | PBS n = 12;  VEGF-A n = 12;  Spike protein n = 6;  EG00229 n = 6;  VEGF-A + EG00229 n = 6;  VEGF-A + Spike protein n = 6; | None |
| Figure 2H | Naïve female rats – paw withdrawal latency: Area over the curve | One-way ANOVA  p <0.0001 | Sidak's multiple comparisons test  PBS vs. VEGF-A165 p=0.0080  PBS vs. Spike p>0.9999  PBS vs. EG00229 p>0.9999  PBS vs. EG00229 + VEGFA165 p=0.1992  PBS vs. VEGFA165 + Spike p>0.9999  VEGF-A165 vs. EG00229 + VEGFA165 p>0.9999  VEGF-A165 vs. VEGFA165 + Spike p0.8178 | PBS n = 12;  VEGF-A n = 12;  Spike protein n = 6;  EG00229 n = 6;  VEGF-A + EG00229 n = 6;  VEGF-A + Spike protein n = 6; |  |
| Figure 3C | Whole cell patch clamp electrophysiology – Peak sodium currents | One-way ANOVA  p = 0.0087 | Holm-Sidak’s multiple comparison post hoc test:  Control (0.1% PBS) vs. VEGFA 1nM p = 0.0247;  Control (0.1% PBS) vs. Spike protein 100nM p = 0.6984;  Control (0.1% PBS) vs. Spike protein + VEGFA p = 0.6272;  VEGFA 1nM vs. Spike protein 100nM p = 0.0034;  VEGFA 1nM vs. Spike protein + VEGFA p = 0.0009;  Spike protein 100nM vs. Spike protein + VEGFA p = 0.7683 | PBS vehicle n = 19;  VEGF-A n = 20;  Spike protein n = 18;  VEGF-A + Spike protein n = 21 | None |
| Figure 3H | Whole cell patch clamp electrophysiology – Peak sodium currents | One-way ANOVA  p = 0.0006 | Dunn’s multiple comparison post hoc test:  PBS vehicle vs. VEGF-A p = 0.0108l;  PBS vehicle vs. VEGF-A + EG00229  p>0.9999;  VEGF-A vs. VEGF-A + EG00229  P = 0.0160 | PBS vehicle n = 12;  VEGF-A n = 12;  EG00229 n = 11;  VEGF-A + EG00229 n = 16 | None |
| Figure 4C | Whole cell patch clamp electrophysiology – Peak N type currents |  | Holm-Sidak’s multiple comparison post hoc test:  Control (0.1% PBS) vs. VEGFA 1nM p = 0.0115;  Control (0.1% PBS) vs. Spike protein 100nM p = 0.9926;  Control (0.1% PBS) vs. Spike protein + VEGFA p = 0.9926;  VEGFA 1nM vs. Spike protein 100nM p = 0.0134;  VEGFA 1nM vs. Spike protein + VEGFA p =0.0353;  Spike protein 100nM vs. Spike protein + VEGFA p = 0.9926 | PBS vehicle n = 20;  VEGF-A n = 15;  Spike protein n = 18;  VEGF-A + Spike protein n = 14 | None |
| Figure 4H | Whole cell patch clamp electrophysiology – Peak N type currents |  | Holm-Sidak’s multiple comparison post hoc test:  Control (0.1% DMSO) vs. VEGFA 1nM p = 0.0021;  Control (0.1% DMSO) vs. Spike protein 100nM p = 0.9898;  Control (0.1% DMSO) vs. Spike protein + VEGFA p = 0.9898;  VEGFA 1nM vs. Spike protein 100nM p = 0.0041;  VEGFA 1nM vs. Spike protein + VEGFA p =0.0050;  Spike protein 100nM vs. Spike protein + VEGFA p = 0.9898 | 0.1% DMSO n = 27;  VEGF-A n = 32;  EG00229 n = 16;  VEGF-A + EG00229 n = 18 | None |
| Figure 5B | Slice electrophysiology – Amplitude of EPSCs | One-way ANOVA  p = 0.1044 | Tukey’s multiple comparison post hoc test:  Control vs. VEGF-A p = 0.9781;  Control vs. VEGF-A + EG00229 p = 0.3770;  Control vs. EG00229 p = 0.9731;  Control vs. VEGF-A + Spike p = 0.2744;  Control vs. Spike p = 0.8701;  VEGF-A vs. VEGF-A + EG00229 p= 0.8322;  VEGF-A vs. VEGF-A + Spike protein p= 0.7486;  VEGF-A + EG00229 vs. VEGF-A + Spike protein p >0.9999 | Control n = 16; VEGF-A n = 14; VEGF-A + EG00229 n = 11;  EG00229 n = 13  VEGF-A + Spike n = 15  Spike n=11 | None |
| Figure 5C | Slice electrophysiology – Frequency of EPSCs | One-way ANOVA  p = 0.0009 | Holm-Sidak’s multiple comparison post hoc test:  Control vs. VEGF-A p =0.0002;  Control vs. VEGF-A + EG00229 p = 0.9582;  Control vs. EG00229 p = 0.9957;  Control vs. VEGF-A + Spike p = 0.8877;  Control vs. Spike p = 0.9966;  VEGF-A vs. VEGF-A + EG00229 p = 0.0012;  VEGF-A vs. VEGF-A + Spike p = 0.0135;  VEGF-A + EG00229 vs. VEGF-A + Spike p = 0.9966 | Control n = 16; VEGF-A n = 14; VEGF-A + EG00229 n = 11;  EG00229 n = 13  VEGF-A + Spike n = 15  Spike n=11 | None |
| Figure 6B | Pre-synaptic fractionation – western blot | Kruskal-Wallis test  P=0.0008 | Dunn's multiple comparisons test  pVEGFR2 :  contra PBS vs. Contra spike p=0.0777  ipsi PBS vs. ipsi spike p=0.0347  contra PBS vs. ipsi PBS p>0.9999  VEGFR2 :  contra PBS vs. Contra spike p=0.6204  ipsi PBS vs. ipsi spike p>0.9999  contra PBS vs. ipsi PBS p>0.9999  NRP1 :  contra PBS vs. Contra spike p>0.9999  ipsi PBS vs. ipsi spike p>0.9999  contra PBS vs. ipsi PBS p>0.9999 |  | None |
| Figure 6C | Spared nerve injury in male rats – paw withdrawal threshold | Two-way ANOVA  p <0.0001 | Dunnet's multiple comparisons test  60 min  PBS vs. Spike 2.14 µg in 5µl p=0.0069  PBS vs. Spike 0.214 µg in 5µl p<0.0001  PBS vs. Spike 0.0214 µg in 5µl p=0.0017  PBS vs. Spike 0.00214 µg in 5µl p>0.9999    120min  PBS vs. Spike 2.14 µg in 5µl p<0.0001  PBS vs. Spike 0.214 µg in 5µl p<0.0001  PBS vs. Spike 0.0214 µg in 5µl p=0.0038  PBS vs. Spike 0.00214 µg in 5µl p>0.9999    180 min  PBS vs. Spike 2.14 µg in 5µl p<0.0001  PBS vs. Spike 0.214 µg in 5µl p<0.0001  PBS vs. Spike 0.0214 µg in 5µl p=0.0951  PBS vs. Spike 0.00214 µg in 5µl p=0.9999    240 min  PBS vs. Spike 2.14 µg in 5µl p=0.0035  PBS vs. Spike 0.214 µg in 5µl p=0.1417  PBS vs. Spike 0.0214 µg in 5µl p=0.993  PBS vs. Spike 0.00214 µg in 5µl p=0.9987    300 min  PBS vs. Spike 2.14 µg in 5µl p=0.0744  PBS vs. Spike 0.214 µg in 5µl p=0.3308  PBS vs. Spike 0.0214 µg in 5µl p=0.9917  PBS vs. Spike 0.00214 µg in 5µl p=0.9984 | PBS n = 12  Spike 2.14 µg in 5µl n = 9  Spike 0.214 µg in 5µl n=10  Spike 0.0214 µg in 5µl n=6  Spike 0.00214 µg in 5µl n=6 | None |
| Figure 6D | Spared nerve injury in male rats – paw withdrawal threshold: Area over the curve | Kruskal-Wallis test | Dunn's multiple comparisons test  PBS vs. Spike 2.14 µg in 5µl p=0.0059  PBS vs. Spike 0.214 µg in 5µl p=0.0019  PBS vs. Spike 0.0214 µg in 5µl p=0.3467  PBS vs. Spike 0.00214 µg in 5µl p>0.9999 | PBS n = 12  Spike 2.14 µg in 5µl n = 9  Spike 0.214 µg in 5µl n=10  Spike 0.0214 µg in 5µl n=6  Spike 0.00214 µg in 5µl n=6 | None |
| Figure 6E | Spared nerve injury in female rats – paw withdrawal threshold | Two-way ANOVA  p <0.0001 | Sidak's multiple comparisons test  PBS vs. Spike 2.14 µg in 5µl  60 min p<0.0001  120 min p<0.0001  180 min p <0.0001  240 min p=0.9798  300 min p>0.9999 | PBS n = 8  Spike 2.14 µg in 5µl n = 7 | None |
| Figure 6F | Spared nerve injury in female rats – paw withdrawal threshold: Area over the curve | Kruskal-Wallis test | Mann Whitney test  PBS vs. Spike 2.14 µg in 5µl: p=0.0003 | PBS n = 8  Spike 2.14 µg in 5µl n = 7 | None |
| Figure 6G | Spared nerve injury – paw withdrawal threshold | Two-way ANOVA  p <0.0001 | Sidak's multiple comparisons test  PBS vs EG00229: time after injection  60 min p=0.0007  120 min p<0.0001  180 min p=0.0009  240 min p=0.0008  300 min p=0.0235 | PBS n = 6  EG00229 n = 5 | None |
| Figure 6H | Spared nerve injury – paw withdrawal threshold: Area over the curve | Mann Whitney test | PBS vs EG00229: p=0.087 | PBS n = 6  EG00229 n = 5 | None |

**Table 1. Gating properties of sodium and calcium currents recorded from DRG neurons^a^**

|  | Sodium | Calcium (CaV2.2) |
| --- | --- | --- |
| Control (0.1%PBS) | |  |
| Activation |  |  |
| *V_1/2_* | -19.9±0.6(19) | -0.5±0.8(20) |
| *k* | 5.5±0.5(19) | 6.1±0.7(20) |
| Inactivation |  |  |
| *V_1/2_* | -42.3±3.9(19) | -20.3±9.1(20) |
| *k* | -14.5±4.0 (19) | -14.8±7.3(20) |
| VEGF-A (1 nM) | | |
| Activation |  |  |
| *V_1/2_* | -22.1±0.4(20) | 0.3±0.7(15) |
| *k* | 4.2±0.4(20) | 5.3±0.6(15) |
| Inactivation |  |  |
| *V_1/2_* | -40.1±2.4(20) | -19.3±5.7(15) |
| *k* | -13.4±2.4 (20) | -14.4±4.7(15) |
| Spike protein (100 nM) | |  |
| Activation |  |  |
| *V_1/2_* | -19.8±0.5(18) | 1.1±0.6(18) |
| *k* | 5.0±0.5(18) | 5.4±0.6(18) |
| Inactivation |  |  |
| *V_1/2_* | -46.4±3.5(18) | -21.1±4.6(18) |
| *k* | -13.1±3.6 (18) | -15.0±4.0(18) |
| VEGF-A (1 nM) + Spike protein (100 nM) | | |
| Activation |  |  |
| *V_1/2_* | -17.1±1.6(19) | -0.8±0.7(14) |
| *k* | 5.7±0.6(19) | 6.0±0.6(14) |
| Inactivation |  |  |
| *V_1/2_* | -45.3±3.3(14) | -24.4±4.5(14) |
| *k* | -13.6±3.2(14) | -13.0±4.0(14) |
| Control (0.1% DMSO) | |  |
| Activation |  |  |
| *V_1/2_* | -19.9±2.2(12) | -2.0±0.6(27) |
| *k* | 6.3±1.4(12) | 5.6±0.6(27) |
| Inactivation |  |  |
| *V_1/2_* | -40.6±2.1(12) | -22.7±4.1(27) |
| *k* | -14.2±3.5(12) | -16.1±3.7(27) |
| VEGF-A (1 nM) | | |
| Activation |  |  |
| *V_1/2_* | -24.8±1.4(12) | 1.5±0.7(32) |
| *k* | 4.3±0.9(12) | 5.6±0.6(32) |
| Inactivation |  |  |
| *V_1/2_* | -42.0±1.6(12) | -22.8±2.8(32) |
| *k* | -11.4±2.3(12) | -13.4±2.8(32) |
| EG00229 (30 μM) | |  |
| Activation |  |  |
| *V_1/2_* | -20.7±1.3(11) | 3.2±0.5(16) |
| *k* | 3.5±1.4(11) | 5.0±0.4(16) |
| Inactivation |  |  |
| *V_1/2_* | -40.9±3.1(11) | -18.1±5.5(16) |
| *k* | -15.1±3.8(11) | -15.9±4.4(16) |
| VEGF-A (1nM) + EG00229 (30 μM) | | |
| Activation |  |  |
| *V_1/2_* | -19.0±1.1(16) | 2.5±0.6(18) |
| *k* | 5.0±1.1(16) | 5.1±0.5(18) |
| Inactivation |  |  |
| *V_1/2_* | -50.2±2.6(16)^b^ | -24.6±4.3(18) |
| *k* | -13.6±2.4(16) | -16.3±4.3(18) |

^a^Values are means ± S.E.M. calculated from fits of the data from the indicated number of individual cells (in parentheses) to the Boltzmann equation; *V_1/2_* midpoint potential (mV) for voltage-dependent activation or inactivation; *k*, slope factor. These values pertain to Fig. 2 of the main manuscript. Only statistically significant differences are indicated within the table. Data were analyzed with one-way ANOVA with Dunnett’s post hoc test.

^b^p=0.0165 comparing Control (0.1% PBS) vs. VEGF-A + EG00229 (one-way ANOVA with Dunnett’s post hoc test)
